# Resolving Symbiodiniaceae diversity across coral microhabitats and reef niches

**DOI:** 10.1101/2024.09.06.611593

**Authors:** Wyatt C. Million, Christian R. Voolstra, Gabriela Perna, Giulia Puntin, Katherine Rowe, Maren Ziegler

## Abstract

Dinoflagellates of the family Symbiodiniaceae are main symbionts of diverse marine animals. A large diversity of Symbiodiniaceae also occur beyond the bounds of their multicellular hosts, occupying environmental niches on coral reefs. The link between spatial diversity at ecosystem scale to microhabitats of Symbiodiniaceae within the coral holobiont are largely unknown. Using ITS2 amplicon sequencing, we compared Symbiodiniaceae communities across four environments (seawater, near-reef and distant sediments, and turf algae mats) and two coral microhabitats (tissue and mucus) on a coral reef in the Red Sea. Analysis of ITS2 sequences revealed that coral and environmental habitats were both dominated by the genera *Symbiodinium*, *Cladocopium*, and *Durusdinium*, but environmental habitats additionally harbored *Fugacium*, *Gerakladium*, and *Halluxium*. Each environmental habitat harbored a distinct Symbiodiniaceae community, with 14-27 % exclusive ITS2 sequences. Nonetheless, 17 ITS2 sequences were shared among all habitat types and were variants defining nearly half of the ITS2 type profiles used to further resolve Symbiodiniaceae identity of coral-based communities. Tissues and surface mucus layers of 49 coral colonies from 17 genera had largely identical Symbiodiniaceae communities. Together with the large difference between environmental Symbiodiniaceae communities and those in the mucus, our results indicate a clear barrier between host-associated and environmental Symbiodiniaceae communities marked by only few shared complete type profiles under normal conditions. It remains to be determined how Symbiodiniaceae community dynamics between coral microhabitats and environmental reservoirs change during coral bleaching events. Monitoring coral colonies after mucus sampling confirmed its suitability for repeated long-term monitoring of coral-associated Symbiodiniaceae communities.

## Introduction

The coral holobiont describes the consortium of eukarya, bacteria, archaea, and viruses that interact through a range of symbioses with their animal host, and each other (1,2). Among the most studied of these relationships is that between the coral and its photosynthetic dinoflagellates of the family Symbiodiniaceae. This symbiosis supports the productivity of the holobiont and the ecosystems they build in otherwise oligotrophic waters (3,4). In exchange for a protective, high nutrient habitat, the algal symbionts can meet the majority of the coral energy requirements (5,6) while the remaining budget can be met through heterotrophy (7–9). Despite this capacity, prolonged or severe disruption of the exchange between members, such as those caused by light, temperature, or nutrient stress can lead to the breakdown of the symbiosis known as coral bleaching (10–12).

The genetic diversity between genera and species within the Symbiodiniaceae is accompanied by functional variation and trade-offs in symbiosis with scleractinian corals (13,14). For example, species within the genus *Cladocopium* may provide more fixed carbon to the host than species within the genus *Durusdinium* (*15*), while associations with members of *Durusdinium* often result in increased thermal tolerance (14,16–18). In addition to these patterns at the genus level, species within each genus and variation within these species present an additional level of functional diversity that can contribute to differences in growth or thermal tolerance of the holobiont (19–22). While coral-Symbiodiniaceae associations tend to be specific, where a coral species is usually associated with a distinct species of Symbiodiniaceae, variation within a host species along environmental gradients has been reported (23,24). For example, the Caribbean coral *Acropora cervicornis* typically exhibits high fidelity with *Symbiodinium fitti* (25,26), while colonies of *Platygyra verwyei* show variation between *Cladocopium* (type C3) and *Durusdinium* (type D1-4)-dominated communities along a temperature gradient (27). Coral-associated Symbiodiniaceae communities within a single colony may also shuffle the relative proportions of existing symbiont genera (28) or strains (19,28) or switch by acquiring new symbionts from external reservoirs, such as seawater, sediments, or neighboring turf algae (29).

Despite the potential for exchange, the Symbiodiniaceae communities found within the host and environmental habitats differ in their diversity, composition, and function (30–32). For example, reef sediment and seawater Symbiodiniaceae taxa are low in abundance in the environment but comprise a large richness and diversity (30,33). In contrast, Symbiodiniaceae cell densities in coral tissues are much higher, but typically less diverse communities with usually one dominant and few low abundant taxa (32). Microhabitats within the coral host, such as the surface mucus layer (SML), also provide opportunities for distinct Symbiodiniaceae communities to contribute to changes in the holobiont that are underexplored to date. The coral SML is a conserved habitat originating from the ectoderm which protects colonies from sedimentation, fouling, and desiccation (34,35). As an environmental interface, the SML also traps and releases free-living and symbiotic microbes thus creating the opportunity for high overlap between microbial communities in the coral SML and environmental habitats (36–38). Despite the microscale distance between the tissue and SML, distinct differences in physical and chemical properties (35,39–41) are able to maintain divergent prokaryotic communities in the two microhabitats (38,42,43).

Previous comparisons of the prokaryotic communities inhabiting the SML and coral tissue showed distinct taxonomic communities (38,42,43) which are likely to be accompanied by functional differences among the microhabitats, such as response to disease (44) or environmental sensitivity (45). However, it is still unknown whether mucus-based Symbiodiniaceae communities mirror those within the coral tissue or if the SML houses distinct symbionts potentially sourced from the environment and/or selectively expelled from the tissue (46). Moreover, identical tissue and mucus communities would suggest that non-destructive assessment of the SML is sufficient as proxy of tissue-bound communities in place of destructive tissue sampling. To elucidate fine scale variation across microhabitats within coral hosts (47,48), we compared Symbiodiniaceae communities between the coral tissue and SML to access the variation among these microhabitats.

Here, we used IT2-type profiles predicted by SymPortal (49) to compare the Symbiodiniaceae communities of the tissue and SML of coral colonies across 17 genera occurring in the central Red Sea. Simultaneous sampling of tissue and mucus was accompanied by environmental sampling of ambient seawater, of sediment under and 2 m away from coral, and of co-occurring turf algae in order to compare coral and environmental Symbiodiniaceae reservoirs. To evaluate the effectiveness of mucus sampling as an minimally invasive alternative to destructive tissue sampling, mucus extraction protocols used in the field were replicated on aquarium corals to track potential damage over time. Significant differences between each environmental reservoir, based on ITS2 sequence variation, suggested distinct communities occupy microhabitats on reefs while the number of sequence variants only returned from non-coral sources highlight the large portion of Symbiodiniaceae sequence diversity found outside the holobiont (although not indicative of Symbiodiniaceae species diversity per se). However, the potential for variation within a Symbiodiniaceae genome precludes the use of ITS2 sequence variants alone to assign this genetic variation to distinct individuals or species present in non-coral habitats where community composition does not meet the assumptions of those *in hospite* (i.e., the presence of only one taxon per Symbiodiniaceae genus). In contrast, the ITS2 type profiles assigned based on the presence and relative abundance of ITS2 sequence variants within the tissue and mucus samples showed that the species-level composition between the two microhabitats are nearly identical suggesting similar communities can be maintained across the drastic physical and chemical clines occurring over small spatial scales. Considering the absence of negative physical consequences in aquarium trials, mucus sampling appears to be an adequate, minimally invasive approach to evaluate coral-associated Symbiodiniaceae diversity.

## Methods

### Sample collection

Fifty coral colonies were sampled to compare the Symbiodiniaceae communities between coral host tissue and mucus *in situ*. In August and September 2016, five survey plots were established on the ocean-facing side of Sha’ab reef, a nearshore reef in the central Saudi Arabian Red Sea (22.20355 N, 39.00035 E). Sha’ab reef has a terrace structure, where the shallow reef flat gently slopes to 3-5 m depth to an edge that is overgrown by coral colonies. Below the corals, a large sandy area of approx. 10 m width extends outward from the edge where the next deeper coral terrace starts. Five plots were established along the first shallow reef edge with a distance of 50 m to each other. At each plot we sampled 10 coral colonies growing next to each other along the reef edge. This resulted in samples from 17 coral genera distributed randomly among the five plots. To avoid artifacts due to species misidentification, colonies were identified through the photographic record to genus level only.

Mucus samples were taken prior to destructive tissue sampling by irritating the surface of the colony with the tip of a 10-ml plastic Pasteur pipette for approximately 10 seconds. This process generated additional mucus that we captured with the pipette. Each pipette containing a sample was bent shut underwater and fixed with a rubber band. Upon return to the boat, all pipettes were opened with the tip facing up and placed on ice for 10 min to let the mucus settle. We then removed the supernatant seawater and a visible mucus sample of approx. 0.5-1 ml was collected from each coral colony into a 2 ml cryovial that was immediately flash-frozen. Host tissue samples were collected on the same spot of each colony as the mucus sample using hammer and chisel. Upon return to the boat each coral fragment was rinsed with filtered seawater to remove surface mucus and immediately flash-frozen. One mucus sample was lost during sample preparation, resulting in 49 sets of paired tissue and mucus samples.

In addition to the two coral-associated microhabitats, we sampled Symbiodiniaceae communities from four distinct environmental reservoirs. We collected five replicate 1-liter samples of seawater per plot from approximately one meter above the sampled coral colonies (n = 25). Seawater samples were immediately stored on ice, upon return to the lab water samples were filtered (0.22 µm) to retain Symbiodiniaceae, and the filters were frozen for DNA extraction. The sediment directly at the reef edge below the sampled coral colonies (near-reef sediment) and 2 m away from the reef (distant sediment) was sampled along each plot (n = 4 per plot, n = 20 per sediment type). For each sample, we gently guided a 50 ml Falcon tube along the sediment surface to fill up the tube with the upper layer of sediment. Sediment samples were flash-frozen on board. Turf algae matts were scraped off dead coral skeletons and reef surfaces with tweezers within 1 m distance from the sampled corals, transported in zip-lock bags, and allowed to drain on board before flash-freezing.

### Sample processing and sequencing

Coral tissue was sprayed off from frozen coral fragments using airflow from a sterile, 1 ml pipette tip connected via a rubber hose to a benchtop air pressure valve with 1 ml of lysis buffer AP1 (Qiagen). Following this, an aliquot of 100 µl of the tissue slurry was added to 300 µl of buffer AP1 in a 1.5 ml tube with 100 µl of glass beads. Samples were then placed in the TissueLyser (Qiagen) for 90 seconds at 30 Hz and then transferred to a new 1.5 ml tube. Coral mucus samples were defrosted at room temperature, pipetted up and down to mix, and 100 µl of mucus was added to 300 µl of Buffer AP1 in a 1.5 ml tube.

Seawater filters were thawed and cut in thirds and one third was transferred to a 1.5 ml tube with 400 µl of Buffer AP1. Turf algae samples were transferred to 2 ml tubes with 1 ml of lysis buffer AP1. After brief vortexing, samples were mixed on a rotating wheel for 30 min and then 400 µl was transferred to a new 1.5 ml tube. Sediment samples were defrosted at room temperature, vortexed and for each sample 10 ml of sediment was transferred to a 50 ml falcon tube and mixed with 20 ml of lysis buffer AP1. Samples were vortexed briefly and then mixed on a HulaMixer for 30 min. Supernatant (400 µl) was transferred to a new 1.5 ml tube.

All sample types were extracted using Qiagen DNeasy Plant Mini Kit (Qiagen, Germany). RNase A (4 µl) was added, samples were vortexed, and incubated for 10 min at 65 °C with tube inversion every 2 min. DNA extractions were then performed according to the manufacturer’s instructions, with a final elution volume of 100 µl, apart from mucus and seawater samples which were eluted in 50 µl. DNA concentrations were measured using a Nanodrop, and all samples were adjusted to 10 ng/μl, apart from the coral mucus (1 ng/μl) and seawater (5 ng/μl).

The ITS2 rDNA region (ITS2) was amplified using the primers SYM_VAR_5.8S2: 5′ (<u>TCGTCGGCAGCGTCAGATGTGTATAAGAGACAG</u>)GAATTGCAGAACTCCGTGAACC 3′ and SYM_VAR_REV: 5′ (<u>GTCTCGTGGGCTCGGAGATGTGTATAAGAGACAG</u>)CGGGTTCWCTTGTYTGACTTCATGC 3′ (50) (Illumina adaptor overhangs underlined). For all samples, triplicate PCRs were performed using 3 μl of DNA, apart from coral samples where 1 μl of DNA was used, and the Qiagen Multiplex PCR kit and a final primer concentration of 0.5 μM in a reaction volume of 10 μl. Thermal cycling conditions were as follows: 95°C for 15 min, followed by 30 cycles (35 cycles for sediment and turf algae samples) of 95°C for 30 s, 56°C for 90 s, 72°C for 30 s, and a final extension cycle at 72°C for 10 min. Then, 5 µl of the PCR was run on a 1% agarose gel to confirm successful amplification. Triplicates for each sample were pooled, and samples were cleaned using ExoProStar 1-step (GE Healthcare). Samples were then indexed using the Nextera XT Index Kit v2 (dual indexes and Illumina sequencing adaptors added), cleaned, and normalized using the SequalPrep Normalization Plate Kit (Invitrogen). The ITS2 libraries were pooled (4 μl per sample) and concentrated using a CentriVap Benchtop Vacuum Concentrator (Labconco). The quality of the library was then assessed using the Agilent High Sensitivity DNA Kit on the Agilent 2100 Bioanalyzer (Agilent Technologies). Quantification was done using Qubit (Qubit dsDNA High Sensitivity Assay Kit; Invitrogen). Sequencing was performed on the Illumina HiSeq platform in the KAUST Bioscience Core Laboratory (BCL).

### Bioinformatics Analysis

ITS2 sequencing data from environmental samples (seawater, turf algae, sediment) were analyzed in SymPortal (49) separately from coral samples (tissue and mucus). Due to the potential high within-genus diversity of Symbiodiniaceae in environmental samples, the assumptions for the full SymPortal analysis could not be met and environmental samples were therefore analyzed as post-MED sequences. Coral samples were analyzed with the full SymPortal workflow which produced post-MED sequences as an intermediate step as well as determined ITS2 type profiles as the final output. Post-MED ITS2 sequence count tables were used for direct comparisons of environmental samples and coral samples, while the ITS2 type profile count tables were used to compare coral tissue and mucus samples. Three turf algae samples with less than 15,000 reads were removed prior to analysis. Additionally, retained low abundance ITS2 sequences (< 0.01 %) across all samples were removed with the purgeOutliers function from the *MCMC.OTU* package in R (51). The 0.01 % threshold represents a conservative cutoff relative to the standard of 0.1 % (52) that will preserve ITS2 sequences indicative of potential free-living Symbiodiniaceae at especially low abundances in the environment.

Differences in Symbiodiniaceae community composition in coral and environmental samples were visualized with a Principal Coordinates Analysis based on Bray-Curtis dissimilarity with the *phyloseq* package (53). The dispersion (i.e., beta diversity) of each Symbiodiniaceae reservoir was compared with an analysis of variance (ANOVA) and post-hoc Permutation test of multivariate homogeneity of groups dispersions in the *vegan* package (54). Differences in the composition between the six reservoirs and sampling plots were assessed with a fully-crossed two-factorial Permutational Multivariate Analysis of Variance (PERMANOVA) in *vegan* (*54*). Separate PERMANOVAs on coral and environmental datasets were also conducted to specifically test for differences between coral mucus and tissue communities and between the environmental reservoirs. Patterns of shared and exclusive ITS2 sequences across the six sources were explored with plots from the *UpSet* package (55). We used the SymPortal ITS2 type profile predictions as proxies for Symbiodiniaceae species and further evaluated potential differences between tissue and mucus reservoirs with this increased taxonomic resolution. All analyses and visualization was completed in the R 4.1.2 environment (56).

### Assessment of tissue integrity following mucus sampling

To identify potential effects of mucus sampling on long-term coral tissue integrity, we simulated the sampling procedure on 2-3 genetically distinct individuals of *Galaxea fascicularis*, *Acropora muricata*, *Porites rus*, *Stylophora pistillata*, and *Pocillopora verrucosa* originating from the Red Sea and Indopacific and compared their recovery to untouched clones. At the Ocean2100 coral aquarium facility (57) at Justus Liebig University Giessen, Germany corals were sampled following the *in situ* procedure, i.e. irritating ~2 cm^2^ of the colony surface with the tip of a plastic Pasteur pipette for 10 seconds. We then tracked 15 disturbed and 15 undisturbed colonies photographically before, immediately after, and 1, 3, 5, 7, and 14 days post-sampling. These were visually inspected and photographically documented to monitor the integrity of the tissue in the sampled area (e.g., appearance of lesions). Pictures were taken with the ONEPLUS A3003 camera (Sony IMX 298 Sensor, 16 MP, 1.12 µm, PDAF, OIS, f/2.0) in the same aquarium where corals were reared and sampled. Camera positions and settings were optimized for each subject and once established remained constant across days.

## Results

### ITS2 sequencing and SymPortal data analysis

ITS2 sequencing with Illumina HiSeq resulted in an average of 220,017 reads per sample after removing three samples with less than 15,000 reads. Post-MED output from SymPortal contained 1,261 unique ITS2 sequences across environmental samples (seawater, turf algae, near-reef sediment, and distant sediment) and 744 unique sequences from coral samples (tissue and mucus). After removing low abundant ITS2 sequences (< 0.01 %, corresponding to 1,240 reads), 411 and 208 unique ITS2 sequences remained in environmental and coral samples, respectively. Together, 534 unique ITS2 sequences were identified from all samples with 195 (36.5 %) belonging to the genus *Symbiodinium*, 254 (47.6 %) to *Cladocopium*, 73 (13.7 %) to *Durusdinium*, 1 (0.1 %) to *Fugacium*, 10 (1.9 %) to *Gerakladium*, and 1 (0.1%) to *Halluxium*.

### Symbiodiniaceae diversity differs among environmental reservoirs

SymPortal identified 411 unique ITS2 sequences in the four types of environmental samples belonging to six Symbiodiniaceae genera. Based on ITS2 sequence diversity, Symbiodiniaceae community composition of these environmental reservoirs was significantly different (PERMANOVA, R^2^ = 0.154, F = 6.76 p = 0.001, Table S1) which was confirmed with the principal component analysis (Figure 1). Communities of both near-reef and distant sediment samples had the largest similarity, but were distinct from each other (F = 2.723, p = 0.003), and significantly different from those in turf algae (F= 8.11, p = 0.001; F = 4.78, p = 0.001, respectively) and seawater (F = 18.26, p = 0.001; F = 9.18, p = 0.001, respectively), which were also significantly different from each other (F = 4.69, p = 0.001, Table S2). Additionally, there was no effect of plot or interaction of sample source and plot on community composition (p = 0.38; p = 0.98, respectively). Overall dissimilarity (i.e., beta diversity) in Symbiodiniaceae composition among samples of a given habitat was significantly lower in seawater (0.366) relative to near-reef sediment (0.468), distant sediment (0.541), turf algae (0.601), SML (0.527), and coral tissue (0.528), which were all statistically similar except for near-reef sediment samples which had significantly lower dissimilarity than turf algae (Table S4).

**Figure 1:**
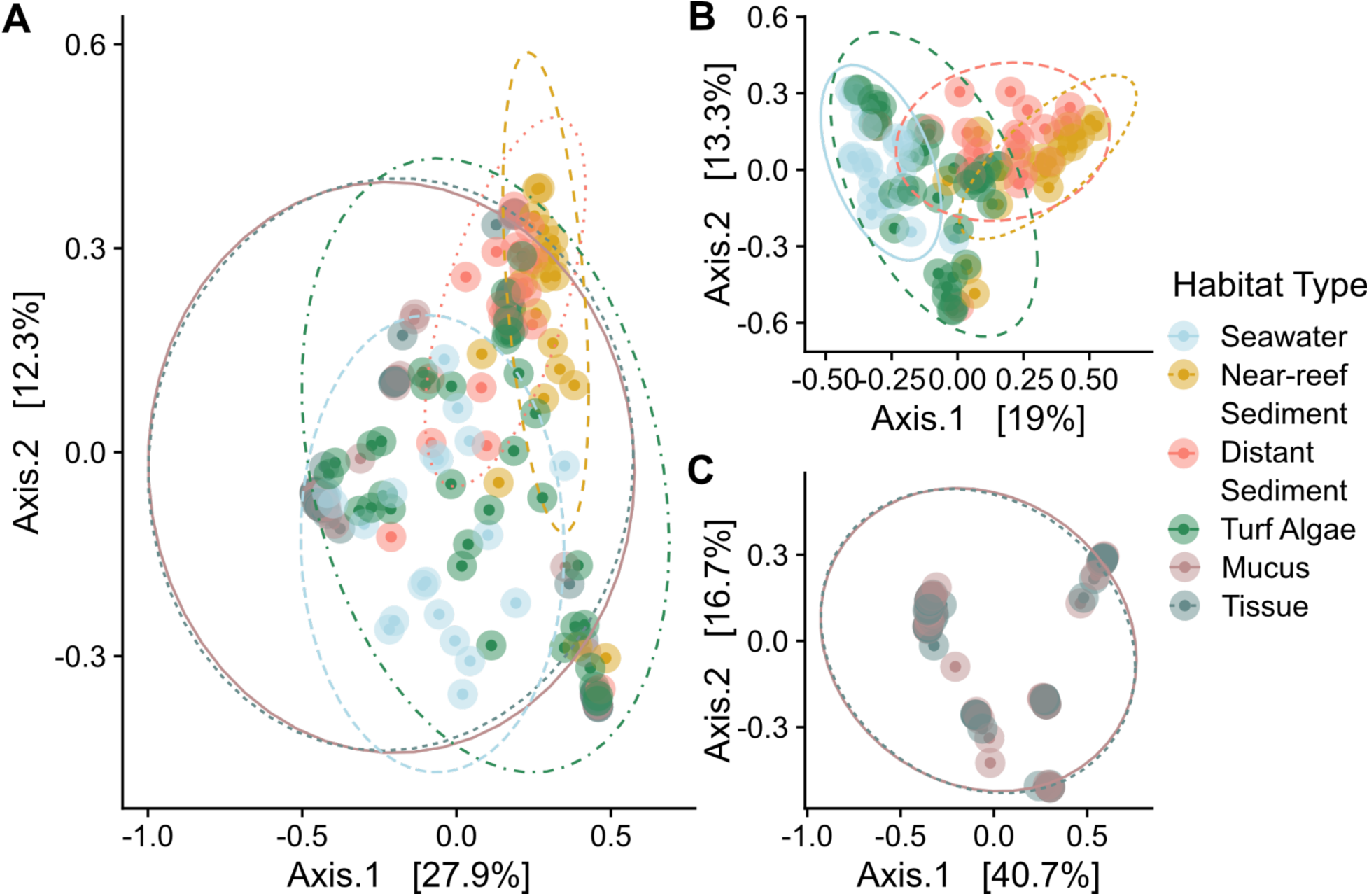
Symbiodiniaceae community composition in environmental reservoirs in a coral reef in the Red Sea based Principal Coordinates Analysis of Bray-Curtis distances of ITS2 sequences for A) all samples, B) samples originating from only environmental sources (i.e., seawater, near-reef sediment, distant sediment, and turf algae), and C) samples collected from only coral sources (i.e., coral tissue and mucus).

The seawater samples were largely dominated (relative abundance > 0.5) by *Cladocopium* spp., though the majority of samples additionally had *Symbiodinium* spp. present at relative abundances of 10 to 65 % (Figure 2A, Figure S1). Near-reef sediment samples were almost exclusively dominated by *Symbiodinium* spp. with *Cladocopium* spp. dominating one sample and co-dominating another (Figure 2C, Figure S1). Symbiodiniaceae communities in distant sediment and turf algae samples showed a larger range of genus-level distribution, occasionally dominated by a single genus, but with most samples displaying more balanced composition relative to near-reef sediment (Fig 2B&C). The highest relative abundances of *Durusdinium* spp. were also found in turf algae and near-reef sediment. Though rarely at high abundance, sequences of *Fugacium* were only seen in sediment habitats, while *Gerakladium* and *Halluxium* sequences were found in seawater and both sediment types.

**Figure 2:**
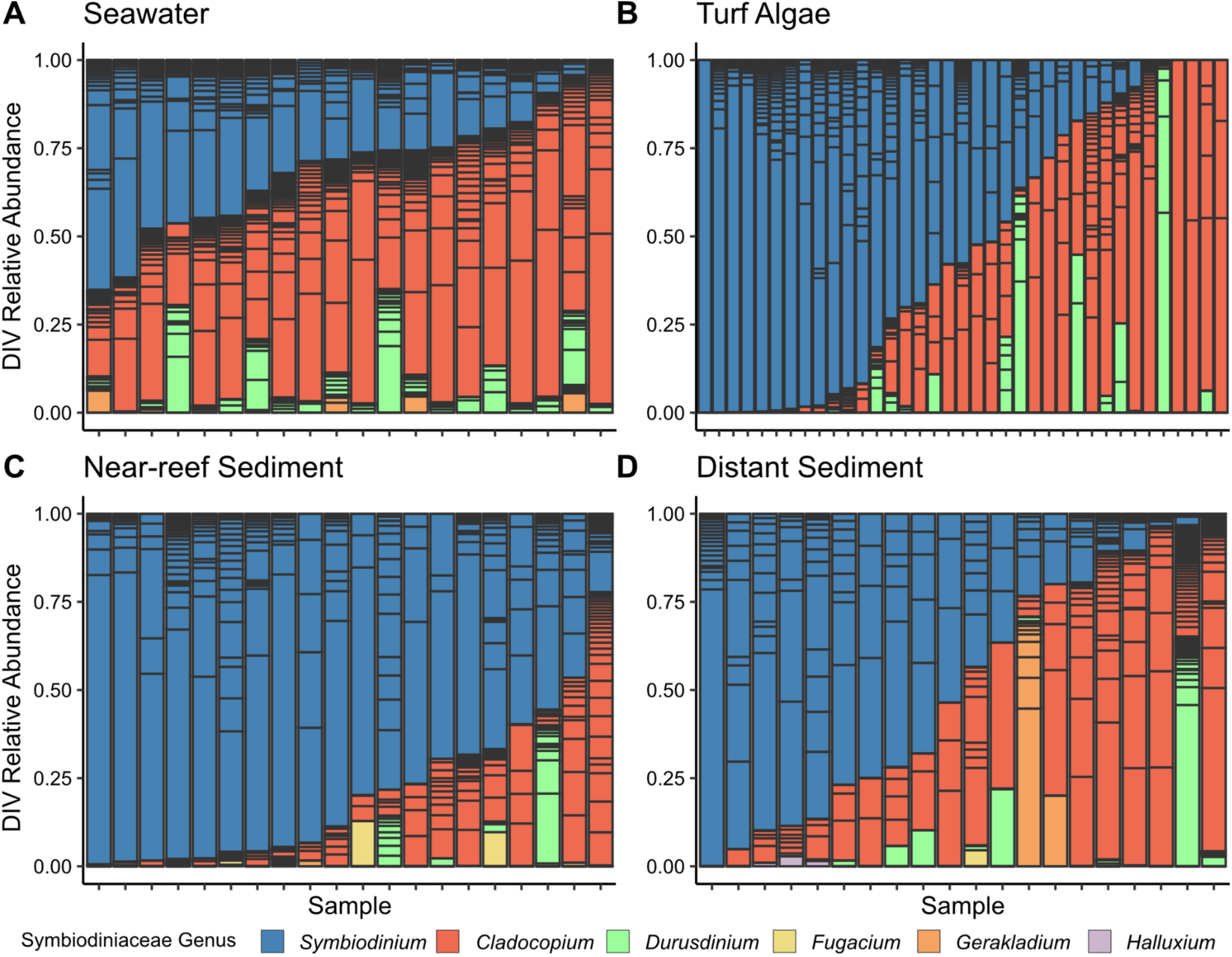
Relative abundance of unique ITS2 sequences in samples from seawater (A), turf algae (B), near-reef sediment (C), and distant sediment (D) environmental sources on a coral reef in the Red Sea. Individual sequences are colored by genus and separated by horizontal bars within each sample. See Figure S3 for a more detailed version colored by individual sequences.

At the individual sequence level, seawater samples had the highest sequence richness overall (229) and average number of distinct sequences per sample (42.95; Figure S2), followed by turf algae (207; 16 per sample), distant sediment (169; 21.2 per sample) and near-reef sediment (140; 21.05 per sample). Only 16 turf algae, 10 near-reef sediment, and 2 distant sediment samples had any sequences occupying greater than 50 % relative abundance and only 12 sequences occupied these dominant roles (Figure S3). Nearly half of the sequences within a habitat type occurred in a single sample only (41-51 % of sequences; Figure S4) and it was rare for sequences to occur in more than five samples. More broadly, the four environmental reservoirs were characterized by large proportions of exclusive sequences only found in a single reservoir (Figure 3). Specifically, 27.5 % of turf algae, 24.4 % of seawater, 13.6 % of near-reef sediment and 26 % of distant sediment sequences were exclusive to their respective reservoirs. Figure 1 suggests seawater and turf algae samples are similar while the two sediment habitats have high overlap. Among the sequences identified in seawater and turf algae, 38 % were shared between the two habitats while 32.6 % of sequences were shared between near-reef and and distant sediment habitats. Only 1 sequence was exclusively shared between all environmental reservoirs, but not corals, and 17 were shared between all environmental reservoirs and the coral samples (Table S6).

**Figure 3:**
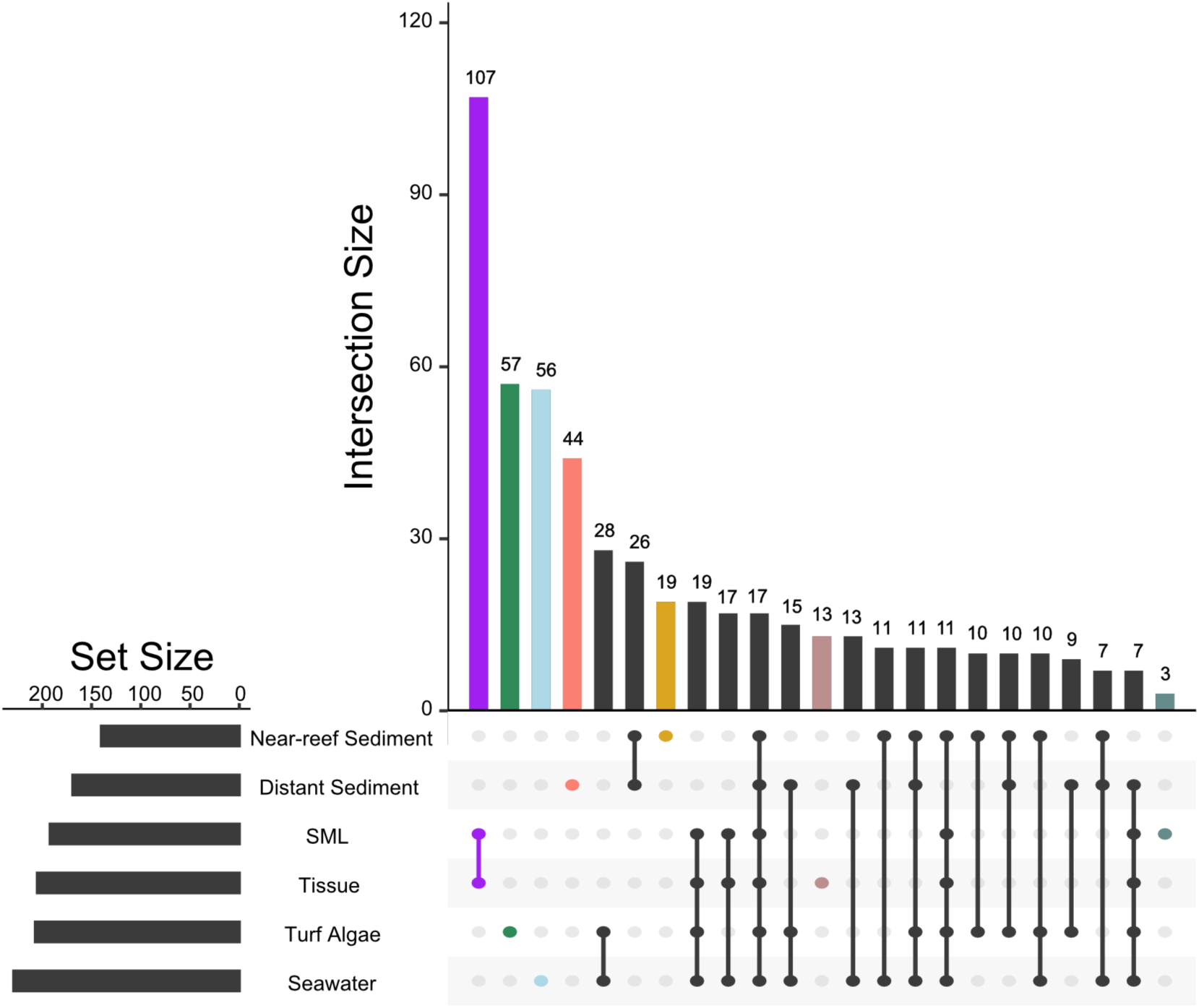
Distribution of shared ITS2 sequences between combinations of coral and environmental sources. The number of sequences shared between a given set of sources (distinguished by filled points) is displayed by the vertical barplot. Total number of unique sequences for each source is shown in the horizontal bar plot. Only combinations with more than 3 shared sequences are shown here (all combinations included in Figure S5).

In agreement with the community ordination, coral samples shared more sequences with turf algae and seawater exclusively, than with sediments (19 vs. 5). Considering that collections of sequences, as opposed to single sequences, represent Symbiodiniaceae individuals due to the potential for intragenomic diversity, we searched for complete sets of sequences in free living samples that correspond to the set of Defining Intragenomic Variants (DIVs) that compose the ITS2 type profiles from coral sources. Of the 41 ITS2 type profiles found in corals, 21 complete DIV sets were present in environmental samples with 20 sets present in seawater, 14 in turf algae, 10 in near-reef sediment, and 7 in distant sediment (Figure S6).

### Coral tissue and mucus harbor nearly identical Symbiodiniaceae communities

Overall Symbiodiniaceae community compositions of coral tissue and mucus samples were indistinguishable from each other when analyzed together with environmental reservoirs (p = 1) and when tested alone (p = 0.99). Similarly, only 19 of 208 sequences found in coral tissue and mucus were not shared between these compartments, and these were all low abundant sequences occurring at < 2 % relative abundance in any given sample apart from one sequence (X1778942_C) occurring in a mucus sample at 3.7%. Nearly half of the sequences were shared between the tissue and mucus exclusively (Figure 3, Figure S5).

To further explore the similarity between tissue and mucus compartments and to help resolve intragenomic variation within the Symbiodiniaceae ITS2 region, we utilized the IT2 type profiles from SymPortal as proxies for genotypes. Based on the 98 coral samples, SymPortal predicted 41 unique ITS2 type profiles belonging to *Symbiodinium* (16 distinct profiles), *Cladocopium* (17), and *Durusdinium* (8), of which 26 occurred as dominant type profiles in samples. Of these ITS2 type profiles, 19 occurred in both the tissue and mucus, 15 in the tissue alone, and 7 in the mucus alone. Interestingly, these 26 dominant type profiles tended to be exclusive to a single coral genus if belonging to *Symbiodinium* or *Durusdinium* while *Cladocopium* type profiles were often dominant across genera apart from four type profiles consisting of C15-related DIVs that were exclusive to *Porites* (Figure S7). When comparing the Symbiodiniaceae community between corresponding tissue and mucus samples per colony, 47 of 49 pairings had identical dominant sequences (i.e., most abundant DIVs within ITS2 type profile), with 33 of the 49 pairings having identical dominant ITS2-type profiles and in two cases, the profile of background types also matched across the compartments (Figure 4). In 16 pairings ITS2-type profiles were not identical, however the most abundant sequence was the same in 14 pairings (Table S7). Of the two non-matching pairs, one set of samples from *Ctenactis* spp. differed in composition of *Cladocopium* sequences and the tissue of a *Pavona* spp. returned a *Durusdinium* ITS2 type profile while its corresponding mucus appeared to be dominated by a *Cladocopium* ITS2 type profile (Figure 4B). As expected, the 16 non-identical pairings showed a similar distribution of ITS2 sequences despite receiving differing ITS2 profile designations from SymPortal (Figure 4B).

**Figure 4:**
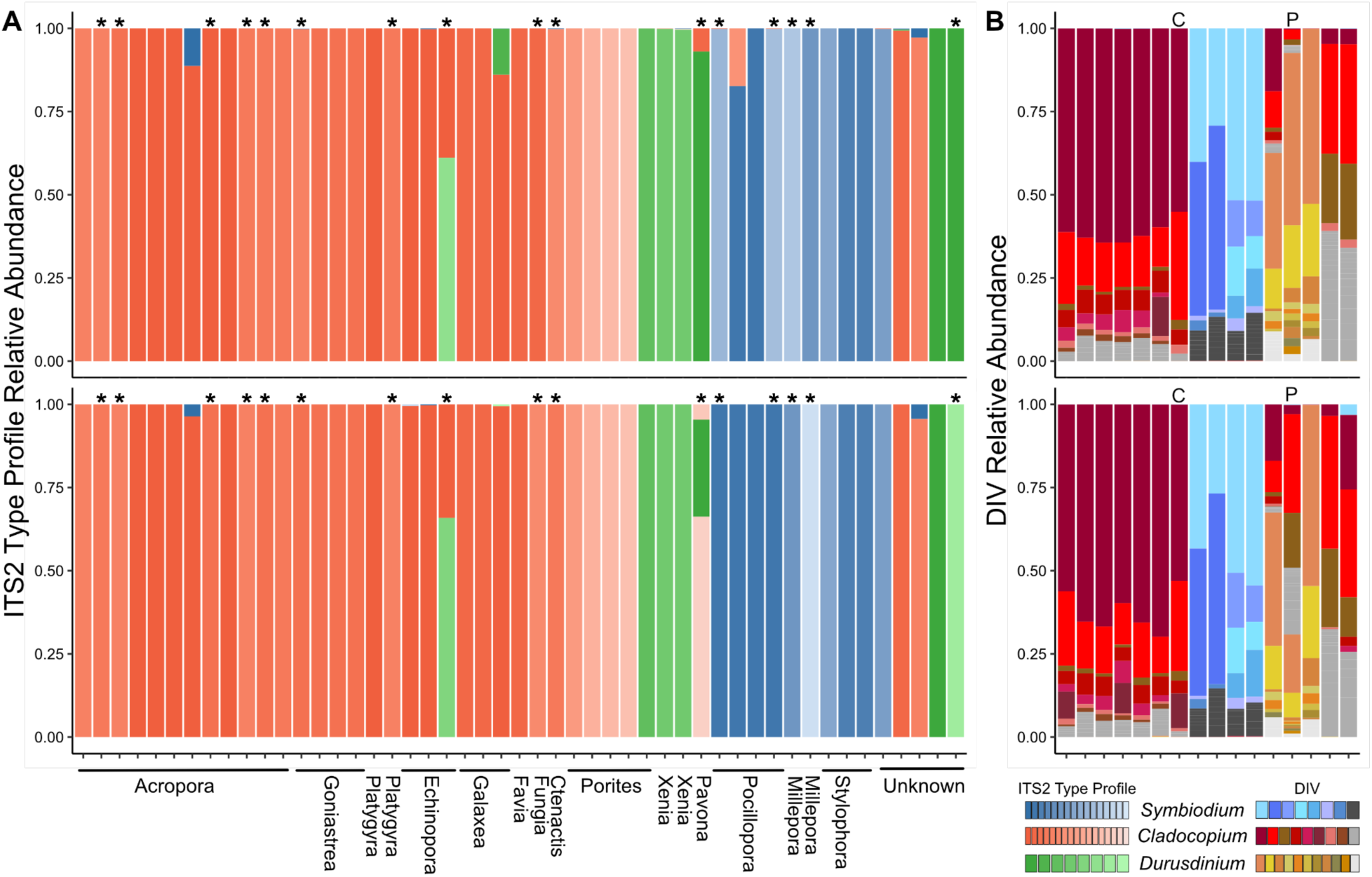
Relative abundance of ITS2 type profiles (A) in coral tissue (top) and SML (bottom). Tissue-SML pairings with non-matching ITS2-type profiles are labeled with an asterisk (*). For these non-matching pairings, the relative abundance of the DIVs within each sample are shown in panel B. Samples in panel B are grouped by DIV community similarity rather than host genus as in panel A. Here, the *Cnetactis* and *Pavona* samples which showed the largest differences in ITS2 DIV relative abundance between tissue and SML are labeled with a C and P according to genus, respectively.

### Mucus sampling is non-invasive

In total 15 coral colonies were sampled and tracked for 14 days post-sampling together with 15 non-treated controls. The only physical effect of the mucus sampling procedure was seen immediately after a colony was sampled and consisted of retreated polyps over a roughly 2 cm^2^ area of tissue (Figure S8). This was more pronounced for species with larger polyps (i.e., *Galaxea fascicularis*). From one day after mucus sampling, none of the colonies showed signs of disturbance or damage, mirroring their untreated clonemates and the pre-sampled condition (Figure S8).

## Discussion

Considering the potential of free-living Symbiodiniaceae communities as reservoirs that offer additional genetic and functional diversity besides that retained in endosymbiotic communities (30,32,33,58), their further documentation will contribute to the growing understanding of the diversity of Symbiodiniaceae. In particular, investigations pairing free-living and *in hospite* communities have the potential to shed light on the exchange between reservoirs. To this end, we explored differences in the Symbiodiniaceae communities of four environmental and two coral-based habitats to identify overlap/segregation that may support potential exchange or isolation of Symbiodiniaceae on natural reefs. We found distinct differences between free-living communities that align with previous reports of Symbiodiniaceae in seawater, sediment, and turf algae niches. Within a coral, however, the nearly identical communities across the tissue and mucus compartments suggest accurate Symbiodiniaceae genotype identification (i.e., ITS2 type profiles) can be made from minimally invasive mucus sampling.

### Distinct free-living Symbiodiniaceae in natural reef reservoirs

Symbiodiniaceae have been found to occupy numerous niches on coral reefs outside of their symbiosis within the coral holobiont (30,32,33,59,60) and the genetic and symbiotic potential of these communities is of particular focus given the opportunity for horizontal transmission of symbionts during juvenile and adults stages (29,61). Here, we found distinct Symbiodiniaceae community assemblages among seawater, turf algae, near-reef sediment, and distant sediment microhabitats that align with previous reports of the propensity for free-living Symbiodiniaceae communities to remain spatially heterogeneous. (30,32,33,59,60). Seawater and turf algae samples tended to be more similar with large portions of *Symbiodinium* and *Cladocopium* and low abundances of *Fugacium*, *Gerakladinium*, and *Halluxium* sequences when compared to sediment sources which had the highest proportions of these genera, similar to previous reports (30,32,62). Interestingly, the near-reef and distant sediment habitat types also displayed distinct communities despite being only two meters from each other. The near-reef sediment samples, taken from directly under the adjacent corals, were almost completely dominated by *Symbiodinium* while distant sediments, sampled in an open patch two meters away from corals, were more variable across samples (Fig. 2). Yet, Symbiodiniaceae communities in both types of sediment samples were equally distinct from those associated with corals, illustrating that Symbiodiniaceae originating from the coral colonies are not preferentially settling in sediments in their direct vicinity. Rather, the coral-derived organic matter and metabolites may create a niche in the near-reef sediment supporting a distinct Symbiodiniaceae community (63). Despite these overall differences across small spatial scales, there was no effect of sampling location (i.e. plot) and the type of environmental niche thus strongly structured the Symbiodiniaceae communities. Dissimilarity among samples was similar across environment types except for seawater, which samples had significantly higher homogeneity. This is perhaps expected considering the physical properties of the water column, a habitat type which was also found to support spatial homogeneity of prokaryotic communities (42).

In line with the metacommunity characterisation of Symbiodiniacaeae dynamics on coral reefs, which suggests distinct habitats are linked through exchange between them (30,64), a number of ITS2 sequences were shared between environmental types in the current study. Specifically, the turf algae samples, which had the most unique sequences, still shared nearly 75% of ITS2 sequences with at least one other environment type. Eighteen sequences were present in all environmental habitats which stands in comparison to the lack of universal ASVs reported by Fujise et al. (32) across sediment, seawater, and coral. The presence of individual sequences across environment types, however, does not mean an equal number Symbiodiniaceae taxa or cells are being exchanged. Here, environmental habitats could only be analyzed to the level of sequences which may present in, but cannot be analyzed as, DIV-based ITS2 type-profiles that SymPortal uses to define taxa. Therefore, the presence of a distinct sequence in two environmental samples may arise through 1) the presence of the same taxon in both samples or 2) two different taxa with the same ITS2 copy within their genomes (49). So while half of the ITS2 type profiles were recovered in at least one free-living sample as complete sets of sequence variants (Figure S6), the inability to resolve inter vs. intragenomic variation in environmental samples limited the ability to determine the amount of exchange of Symbiodiniaceae individuals between the habitats explored here.

Conversely, the absence of a particular ITS2 sequence variant from an environmental or coral habitat is a useful proxy for identifying exclusive genomic diversity. For instance, over 300 sequences are only found in the four environmental habitats and while these sequences likely do not represent 300 Symbiodiniaceae genotypes, they do offer genetic diversity that is potentially novel from the coral holobiont. Characterizing these free-living communities is therefore important when considering uptake from environmental reservoirs during bleaching recovery, which may offer the opportunity to acquire new, potentially advantageous symbionts (65,66). Similarly, seeding of juvenile corals by the sediment-aided horizontal transmission of symbionts offer yet another mechanism utilizing the diversity found within reef habitats (61,67),

### Coral tissue and mucus have mirroring Symbiodiniaceae communities

While there have been more frequent empirical comparisons among *in hospite* and various free-living Symbiodiniaceae communities (30,32,33), comparisons between the tissue and SML within the coral holobiont have largely been limited to the prokaryotic communities (38,42,43,68). These studies uncovered that distinct prokaryotic communities persist between coral skeleton, tissue, and SML (38,42), which is in line with the dramatically different environmental conditions across these microhabitats. Similarly for Symbiodiniaceae, the highly regulated symbiosome (40,41) creates a stable environment for intracellular Symbiodiniaceae, while the abiotic conditions of the SML are temporally and environmentally variable and directly interact with seawater (69). Despite these potential barriers, we found indistinguishable communities, high sequence overlap, and preservation of ITS2 type profiles between the coral tissue and SML that suggest abundant and unhindered exchange between the two microhabitats within the holobiont and a clear delimitation of the mucus from the surrounding seawater.

Characterisation of the Symbiodiniaceae in the coral SML is limited and as a result the origins, residence time, and ultimate fate of this community is still unclear. Given the rapid turnover rate of the SML (36), the relatively stable intracellular Symbiodiniaceae communities likely act as a source that repopulates the SML through expulsion of symbionts. Symbionts expelled regularly and while under stress (70,71) likely interact with the SML upon leaving the polyp providing an opportunity to occupy the new niche. While coral often expel degraded Symbiodiniaceae cells, release of healthy individuals (72,73) increases the potential for the formation of an active SML population. Notably, the large overlap in communities between tissue and SML persisted across 17 coral genera, the turnover time of SML Symbiodiniaceae communities remains unclear. Variance among microhabitat-specific prokaryotic communities has been documented to change with environments (74) and considering the potential for selective expulsion (46) and variation in Symbiodiniaceae physiology (13,14), exchange of symbionts between tissue and the SML may also vary over time. Given that 90 % of the sequences found in the tissue were also present in the mucus (Fig. 3), there appears to be a limited degree of selection during exchange between compartments during the time of sampling. Similarly, the maintenance of ITS2 type profiles between tissue and mucus samples confirms the relative abundance of the shared sequences is also preserved across the microhabitat partitions. The coral used here were from an observably healthy reef, lacking disease and bleaching, and therefore offer a baseline condition for additional microbial scale spatial analyses of the coral holobiont (47). While a large taxonomic range of Red Sea corals was investigated here, whether tissue and SML Symbiodiniaceae communities remain homogenous under stress conditions should be addressed with future studies.

The use of the ITS2 type profiles predicted by SymPortal offers enhanced biological resolution when characterizing intergenomic and intragenomic variation within Symbiodiniaceae communities of the tissue and SML. However, of the 49 tissue-mucus sample pairings, 16 returned nonmatching ITS2 type profiles despite the majority of DIVs being present in both microhabitats of 48 pairings (Fig. 4B). Only a single pairing from a *Pavona* colony returned ITS2 type profiles from different genera (*Cladocopium* in tissue, *Durusdinium* in mucus). The remaining mismatched pairings showed less divergence and differed in 1) whether the most abundant DIV in one sample was the most abundant or in co-majority with the second most abundant DIV in the other sample and/or 2) differences in the presence/absence of background (less abundant) DIVs. Given a coral is often dominated by only a single Symbiodiniaceae taxon (49,75–77), it may be unlikely that two highly related but distinct ITS2 type profiles dominate the tissue and SML. Therefore, if the SML community is derived from the tissue, e.g. via discharge from the tissue, the mismatches seen in 15 of 16 pairing likely do not represent true intraspecific variation of Symbiodiniaceae between the compartments.

Beyond providing insights into the fine-scale spatial distribution of Symbiodiniaceae, the similarities seen here highlight the ability of minimally invasive mucus sampling to replace destructive sampling of coral symbionts. About two-thirds of the ITS2 type profiles corresponded between tissue and mucus and 98 % of mucus samples (i.e., all but one) returned the same majority DIV of the tissue and had near-identical type profiles. Depending on the research question and the biological resolution needed, mucus sampling may thus present an alternative and easy sampling method. Currently, the standard procedures for assessing Symbiodiniaceae communities include sacrificing coral tissue to extract symbiont DNA (30,78–81) which may be unavoidable when conducting other phenotypic measures but inhibits repeat sampling when biological material is limited. Here, the SML was sampled *in situ* following protocols commonly used to obtain mucus for assessments of prokaryotic communities (42,82), which were sufficient enough here to return mucus samples distinct from the seawater - a source of potential error (38). Similarly, Symbiodiniaceae DNA was obtained from tissue samples following the common airbrushing method (78) after mucus was removed prior to sample preservation. Therefore, the comparisons between tissue and mucus Symbiodiniaceae communities likely represent accurate biological similarities rather than artificial overlap due to cross contamination. Moreover, the large diversity of coral genera sampled here suggest the homogeneity among tissue and the SML symbiont communities are not taxon-specific, but instead this approach can be applied across coral species. Finally, SML sampling of coral completed under controlled aquarium conditions revealed no short- or long-term harmful effects on colonies, confirming this widely used technique is appropriate for repeat sampling.

### Conclusions

The potential of reef habitats to harbor genetic, functional, and symbiotic diversity is of particular concern given the reliance of scleractinian coral on uptake of exogenous Symbiodiniaceae from environmental pools during the establishment and recovery of the holobiont. Here, the distinct, yet overlapping communities that reside within the sediment, ambient seawater, and on neighboring algae occupying Red Sea reefs add to a growing body of evidence that free-living habitats support a metacommunity of Symbiodiniaceae linked with the coral holobiont. This niche partitioning appears to break down across microhabitats within the holobiont, however, with Symbiodiniaceae communities in the coral tissue and SML remaining nearly identical despite persistent differences in the prokaryotic communities across this same scale. While the dynamics of endosymbiotic Symbiodiniaceae communities has been of particular focus, the origin and fate of SML communities may offer additional insight into the maintenance, breakdown, and/or recovery of this ecological important symbiosis. Considering its ease and minimal invasiveness, sampling the SML offers a useful tool to not only investigate coral mucus itself but also to obtain a glimpse of the endosymbiosis within.

## Acknowledgements

We would like to thank Benjamin C.C. Hume for running the data through the SymPortal pipeline and discussions of the results. We acknowledge the KAUST Bioscience Core Lab for assistance with sequencing. Research reported in this publication was supported by baseline funds to C.R.V. from KAUST and funded by the Deutsche Forschungsgemeinschaft (DFG, German Research Foundation) – Project number 469364832 (M.Z.) – SPP 2299/Project number 441832482. The time-series mucus sampling was conducted in the ‘Ocean2100’ global change simulation facility of the Colombian-German Center of Excellence in Marine Sciences (CEMarin) funded by the German Academic Exchange Service.

## Author contributions

M.Z. designed and conceived the experiment; M.Z., K.R., and G.Pu. collected data; G.Pe. generated molecular data; M.Z. and C.R.V. provided reagents/tools; W.M., G.Pu., B.H., C.R.V, and M.Z. analyzed the data; W.M. wrote the manuscript which M.Z., C.R.V. edited; all authors read and approved the final manuscript.

## Data availability statement

Sequence data determined in this study are available under NCBI BioProject ID PRXXXX and included in the SymPortal Database. Source data underlying figures and statistical analyses are provided at https://github.com/wyattmillion/MucusTissueEnvironment_SymITS2.

## Supplementary Material

**Table S1:** Summary of results of the PERMANOVA (adonis2) comparing overall community composition among environment types (seawater, near-reef and distant sediment, turf algae, SML, and tissue) based on Bray Curtis Dissimilarity.

| Factor | DF | Sum of Squares | R2 | F | Pr (>F) |
| --- | --- | --- | --- | --- | --- |
| environment type | 5 | 10.947 | 0.15392 | 6.760 | 0.001 |
| plot | 4 | 1.342 | 0.01887 | 1.036 | 0.376 |
| environment type × plot | 20 | 5.069 | 0.07127 | 0.783 | 0.982 |
| Residual | 166 | 53.764 | 0.75594 |  |  |
| Total | 195 | 71.122 | 1 |  |  |

**Table S2:** Pairwise adonis comparing overall ITS2 community based on Bray Curtis Dissimilarity of each habitat. Adjusted p-value represents Bonferroni & Hochberg multiple test corrected p-value.

| Pairs | F.Model | R2 | P-value | Adjusted P-value |
| --- | --- | --- | --- | --- |
| SML vs tissue | 0.06610665 | 0.00068105 | 1 | 1 |
| SML vs turf algae | 4.43616687 | 0.05016236 | 0.002 | 0.002307692 |
| SML vs distant sediment | 8.32493016 | 0.11052025 | 0.001 | 0.001666667 |
| SML vs near-reef sediment | 15.6134325 | 0.18899387 | 0.001 | 0.001666667 |
| SML vs seawater | 5.1741059 | 0.07168923 | 0.002 | 0.002307692 |
| tissue vs turf algae | 4.5182762 | 0.05047323 | 0.002 | 0.002307692 |
| tissue vs distant sediment | 8.33755081 | 0.10921952 | 0.001 | 0.001666667 |
| tissue vs near-reef sediment | 15.6362111 | 0.18695504 | 0.001 | 0.001666667 |
| tissue vs seawater | 5.29110851 | 0.07219305 | 0.002 | 0.002307692 |
| turf algae vs distant sediment | 4.78247398 | 0.07999793 | 0.001 | 0.001666667 |
| turf algae vs near-reef sediment | 8.11487506 | 0.1285731 | 0.001 | 0.001666667 |
| turf algae vs seawater | 4.69162905 | 0.07859777 | 0.001 | 0.001666667 |
| distant sediment vs near-reef sediment | 2.72306359 | 0.06686785 | 0.003 | 0.003214286 |
| distant sediment vs seawater | 9.17824191 | 0.19454396 | 0.001 | 0.001666667 |
| near-reef sediment vs seawater | 18.258485 | 0.32454633 | 0.001 | 0.001666667 |

**Table S3:** Summary of ANOVA (*vegan*) comparing the homogeneity of habitat type variances (i.e. dissimilarity of samples within a given compartment) based on Bray Curtis Dissimilarity.

|  | **Df** | **Sum of Squares** | **Mean Squares** | **F value** | **Pr (>F)** |
| --- | --- | --- | --- | --- | --- |
| Groups | 5 | 0.795 | 0.159006 | 2.9377 | 0.01404 |
| Residuals | 190 | 10.284 | 0.054125 |  |  |

**Tables S4:** Summary of permutation test for homogeneity of multivariate dispersions for all pairwise comparisons of habitat types. Observed p-values are below the diagonal, permuted p-values are above the diagonal. Average distance to the median, as calculated by betadisper in *vegan*, for each habitat is included in parentheses.

|  | **Seawater** | **Neer-reef**<br>**Sediment** | **Distant**<br>**Sediment** | **SML** | **Tissue** | **Turf algae** |
| --- | --- | --- | --- | --- | --- | --- |
| **Seawater (0.366)** |  | 0.025 | 0.001 | 0.022 | 0.028 | 0.001 |
| **Near-reef Sediment (0.463)** | 0.023 |  | 0.101 | 0.356 | 0.356 | 0.001 |
| **Distant Sediment (0.541)** | 0.000 | 0.114 |  | 0.846 | 0.853 | 0.080 |
| **SML (0.5266)** | 0.023 | 0.372 | 0.844 |  | 0.982 | 0.171 |
| **Tissue (0.528)** | 0.024 | 0.367 | 0.862 | 0.980 |  | 0.184 |
| **Turf algae (0.601)** | 0.000 | 0.000 | 0.080 | 0.150 | 0.164 |  |

**Table S5:** Number of unique DIVs (recorded in Symportal database) present in each habitat type and the average found in any given sample as well as the distribution of a habitat’s DIVs across Symbiodiniaceae genera.

|  | **No. of**<br>**unique**<br>**DIVs** | **Average per**<br>**sample** | *Sym.* | *Clad.* | *Dur.* | *Fug.* | *Ger.* | *Hall.* |
| --- | --- | --- | --- | --- | --- | --- | --- | --- |
| seawater | 229 | 42.95 | 90 | 97 | 31 | 0 | 10 | 1 |
| turf algae | 207 | 16 | 115 | 74 | 18 | 0 | 0 | 0 |
| near-reef sediment | 140 | 21.2 | 71 | 49 | 17 | 1 | 1 | 1 |
| distant sediment | 169 | 21.05 | 57 | 78 | 24 | 1 | 8 | 1 |

**Table S6:** List of ITS2 sequences shared between all free-living and coral habitats.

|  |  |  |
| --- | --- | --- |
| A1 | D3a | X56421_A |
| C1 | D4 | X56861_D |
| C15 | D4c | X8111_A |
| C1b | X4185_A | X8182_A |
| C41 | X4935_D | X9354_A |
| D1 | X56419_A |  |

**Table S7:** List of non-matching tissue and mucus samples and their corresponding dominant ITS2 type profile.

| Source | Coral # | Coral Genus | Sample.ID | Dominant Type Profile |
| --- | --- | --- | --- | --- |
| coral | 21 | Acropora | Coral_ITS_021 | C41-C1-C41b-C41j-C41g-C41w |
| mucus | 21 | Acropora | Mucus_ITS_041 | C41/C1-C41b-C41j-C1b |
| coral | 31 | Acropora | Coral_ITS_031 | C41/C1-C41b-C1b-C41o |
| mucus | 31 | Acropora | Mucus_ITS_050 | C41-C1-C41x-C41b-C1b |
| coral | 38 | Acropora | Coral_ITS_038 | C41/C1-C41b-C41j-C1b |
| mucus | 38 | Acropora | Mucus_ITS_057 | C41-C1-C41b-C41j-C41g-C41w |
| coral | 40 | Acropora | Coral_ITS_040 | C41/C1-C41b-C41j-C1b |
| mucus | 40 | Acropora | Mucus_ITS_059 | C41-C1-C41x-C41b-C1b |
| coral | 46 | Acropora | Coral_ITS_046 | C41-C1-C41x-C41b-C1b |
| mucus | 46 | Acropora | Mucus_ITS_065 | C41-C1-C41b-C41j-C41g-C41w |
| coral | 36 | Goniastrea | Coral_ITS_036 | C41-C1-C41b-C41j-C41g-C41w |
| mucus | 36 | Goniastrea | Mucus_ITS_055 | C41-C1-C41x-C41b-C1b |
| coral | 4 | Platygyra | Coral_ITS_004 | C41/C1-C41b-C41j-C1b |
| mucus | 4 | Platygyra | Mucus_ITS_024 | C41-C1-C41b-C41j-C41g-C41w |
| coral | 8 | Echinopora | Coral_ITS_008 | D1-D4-D17d-D4c-D17e-D6-D1r-D17c-D17j |
| mucus | 8 | Echinopora | Mucus_ITS_028 | D1-D4c-D4-D6-D1h |
| coral | 50 | Fungia | Coral_ITS_050 | C1/C39-C1b-C41-C1ae-C41f-C41a-C1eb |
| mucus | 50 | Fungia | Mucus_ITS_069 | C1/C41e-C1b-C41f-C41a-C41-C39-C1ae-C1f-C1eb |
| coral | 25 | Ctenactis | Coral_ITS_025 | C1/C41e-C1b-C41f-C41a-C41-C39-C1ae-C1f-C1eb |
| mucus | 25 | Ctenactis | Mucus_ITS_045 | C41/C1-C41b-C41j-C1b |
| coral | 23 | Pavona | Coral_ITS_023 | D1-D4-D4c-D1h-D1c-D2 |
| mucus | 23 | Pavona | Mucus_ITS_043 | C1-C1b-C39-C41-C41f-C41a-C1ae-C41e-C1af-C1f-C39a |
| coral | 12 | Pocillopora | Coral_ITS_012 | A1/A1c-A1h-A1cc-A1ce-A1ch |
| mucus | 12 | Pocillopora | Mucus_ITS_032 | A1-A1cc-A1c-A1h-A1q-A1i |
| coral | 9 | Pocillopora | Coral_ITS_009 | A1-A1c-A1cc-A1h-A1bv-A1cb-A1ce |
| mucus | 9 | Pocillopora | Mucus_ITS_029 | A1-A1cc-A1c-A1h-A1q-A1i |
| coral | 17 | Millepora | Coral_ITS_017 | A1/A1k-A1g-A1b |
| mucus | 17 | Millepora | Mucus_ITS_037 | A1k/A1-A1ea |
| coral | 22 | Millepora | Coral_ITS_022 | A1k/A1-A1ea |
| mucus | 22 | Millepora | Mucus_ITS_042 | A1/A1k-A1b-A1z |
| coral | 20 | Unknown | Coral_ITS_020 | D1-D4-D4c-D1h-D1c-D2 |
| mucus | 20 | Unknown | Mucus_ITS_040 | D1-D4-D4c-D4f-D1b-D1h |

**Figure S1:**
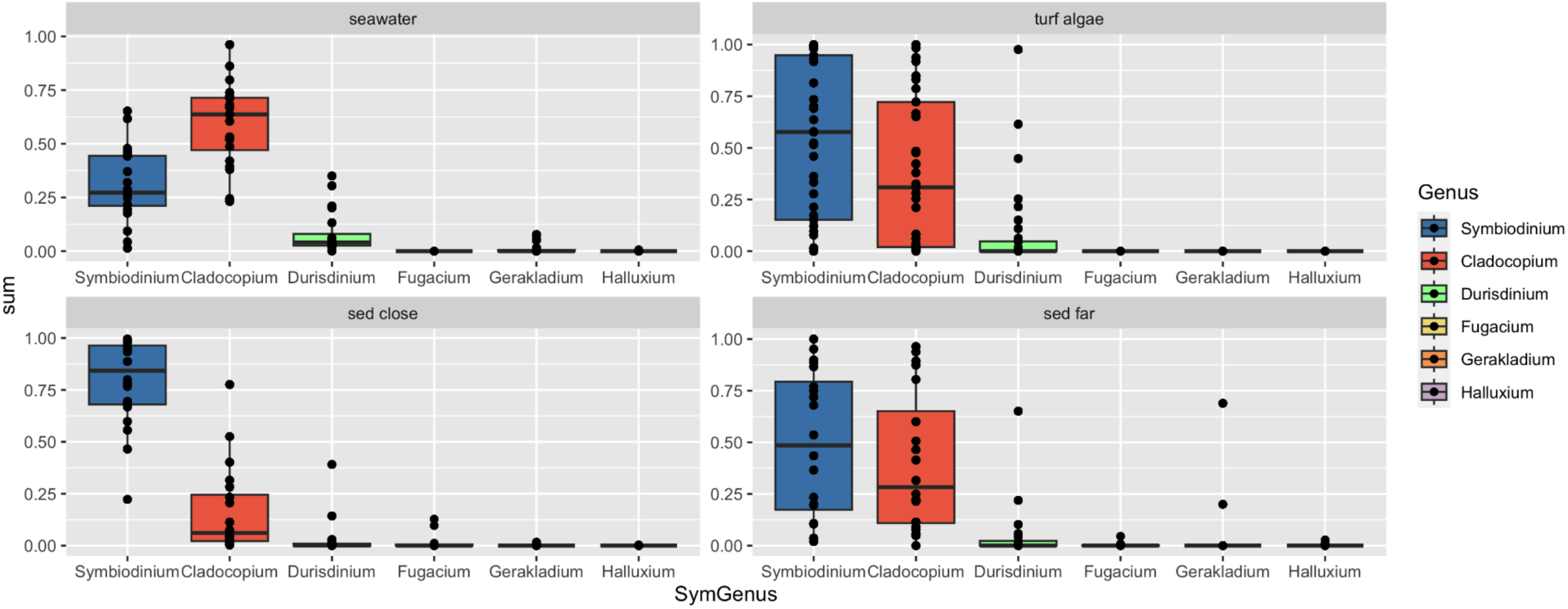
Distribution of genus-level relative abundance within samples from each free-living habitat.

**Figure S2:**
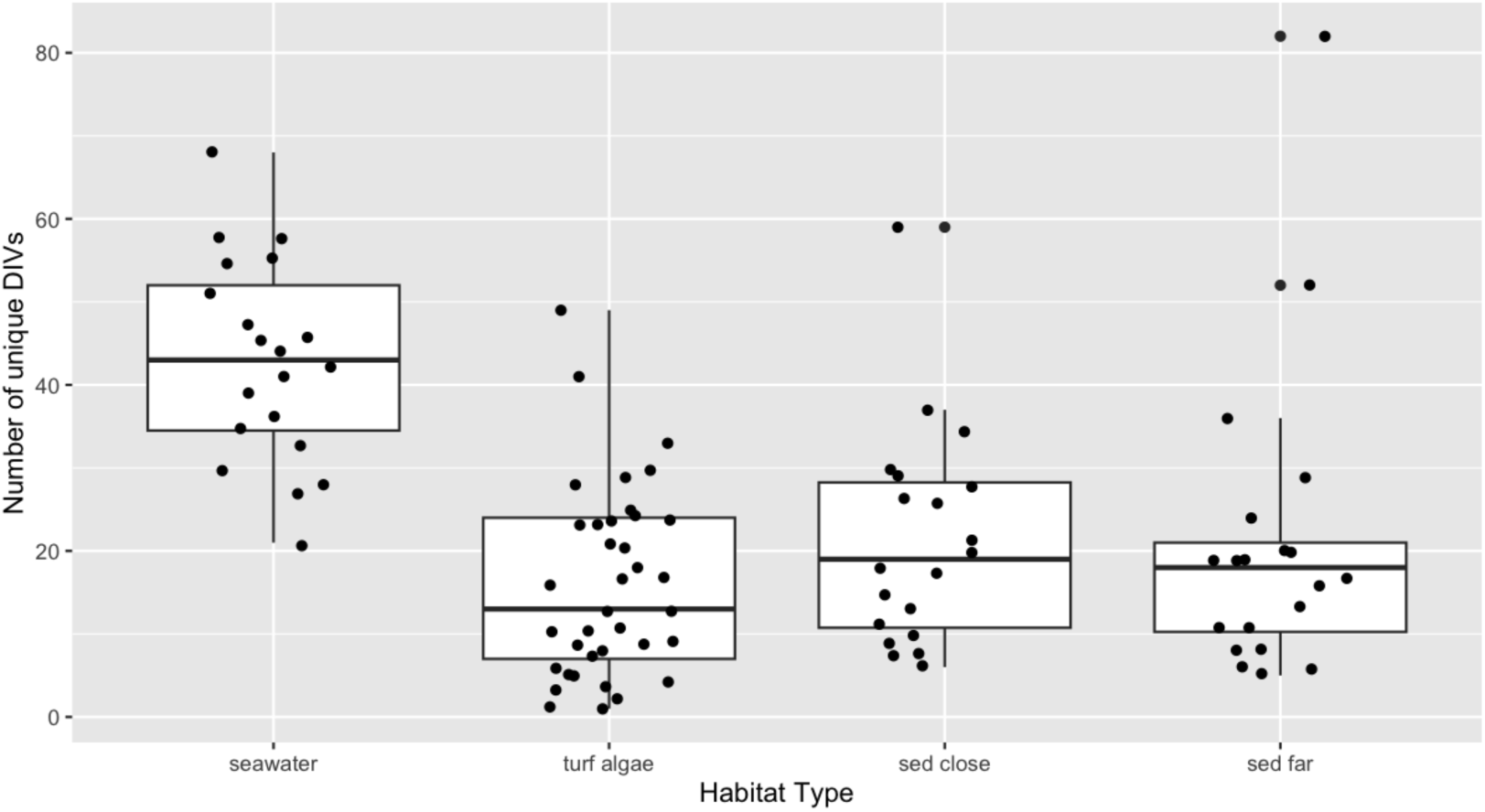
Distribution of unique DIVs per sample within each free-living habitat type

**Figure S3:**
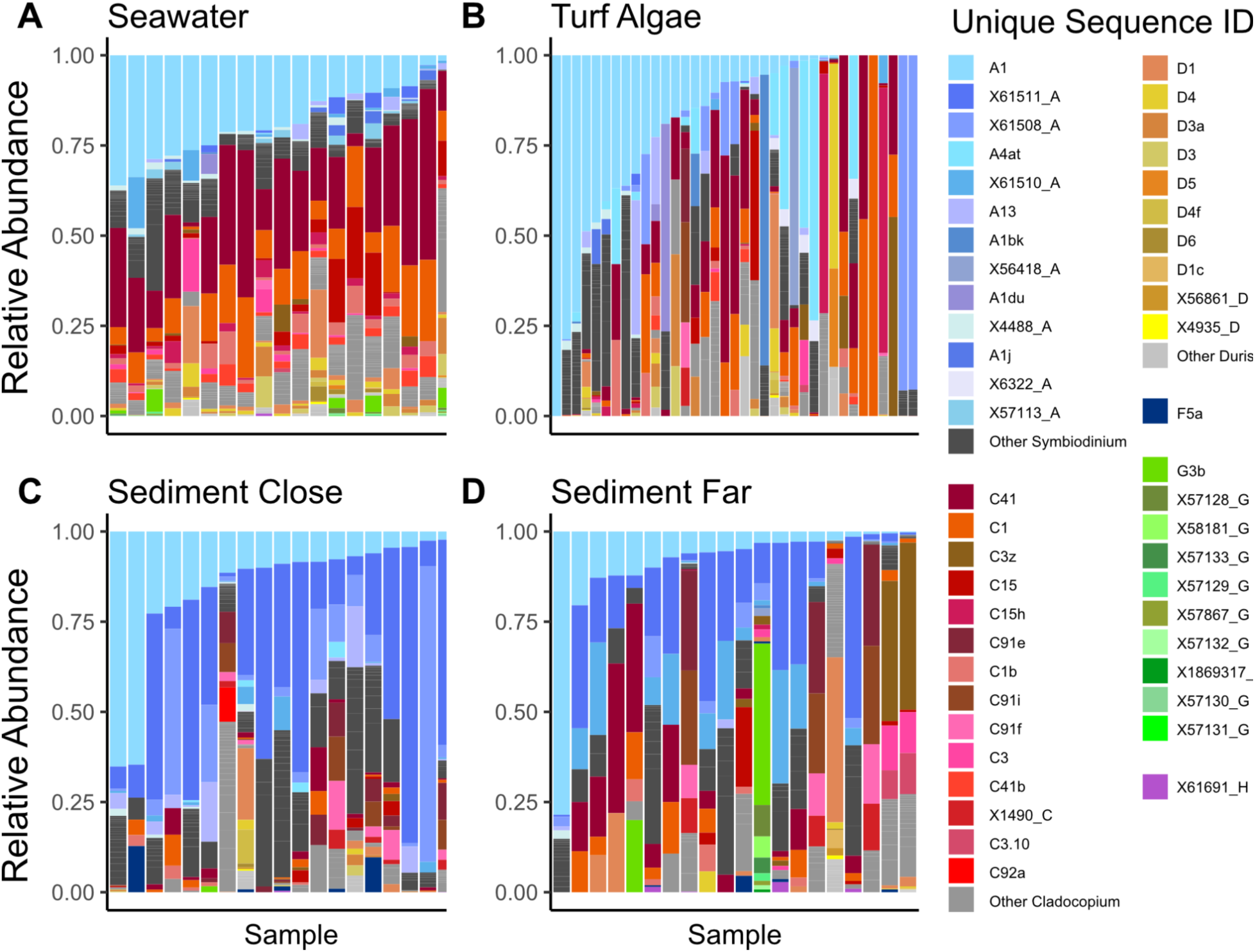
Relative abundance of unique DIVs in samples from (A) seawater (A), turf algae (B), near-reef sediment (C), and distant sediment (D) environmental sources. Sequences representing less than 1% of the total abundance of sequences per Symbiodiniaceae genus across all samples are grouped as “Other”. Two sequences, A13 and A1du, were highly abundant in single samples despite being grouped as “Other Symbiodinium” so these sequences were given a distinct label.

**Figure S4:**
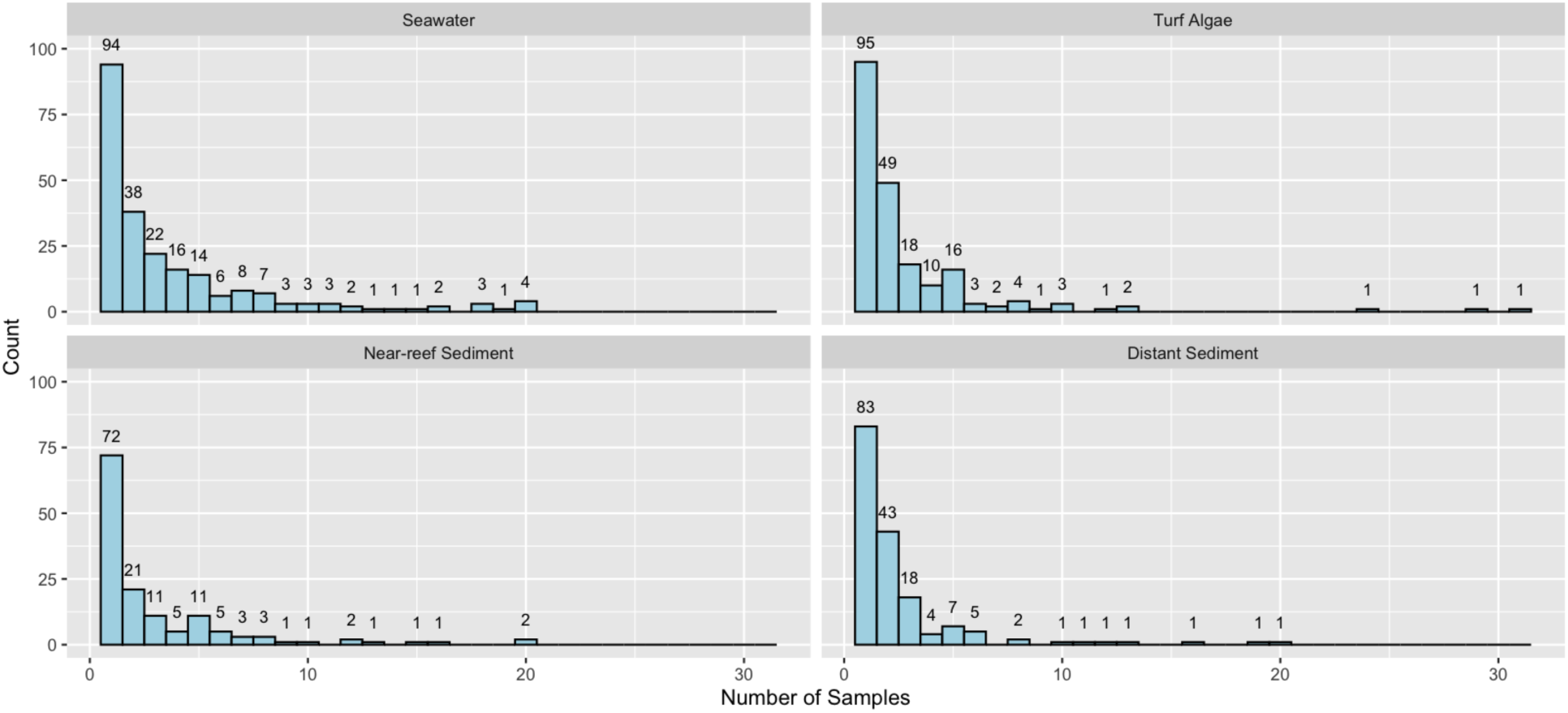
Histogram showing the frequency distribution of distinct DIVs for a given number of samples.

**Figure S5:**
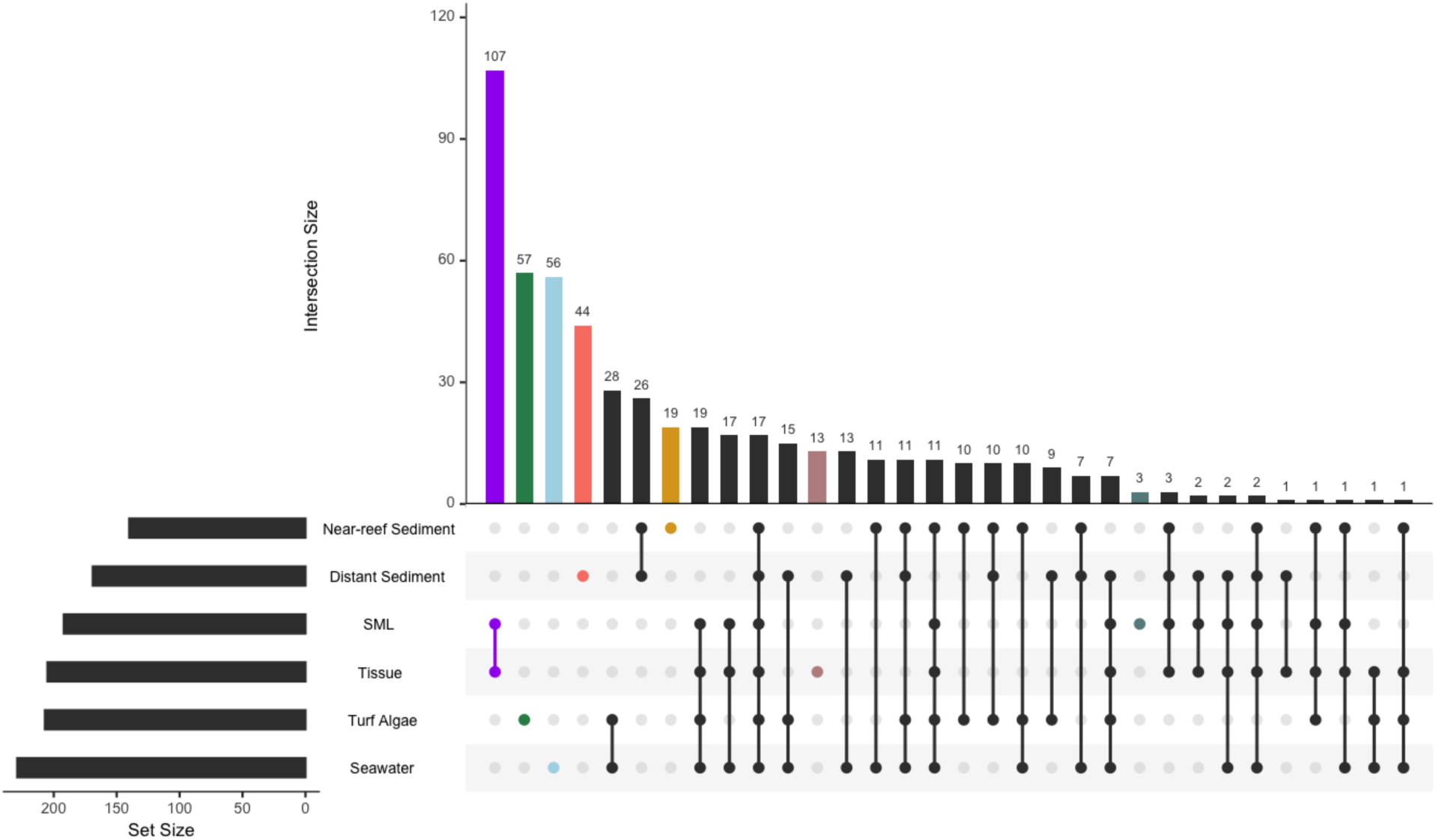
Distribution of shared ITS2 sequences between combinations of coral and non-coral sources. The number of sequences shared between a given set of sources (distinguished by filled points) is displayed by the vertical barplot. Total number of unique sequences for each source is shown in the horizontal bar plot.

**Figure S6:**
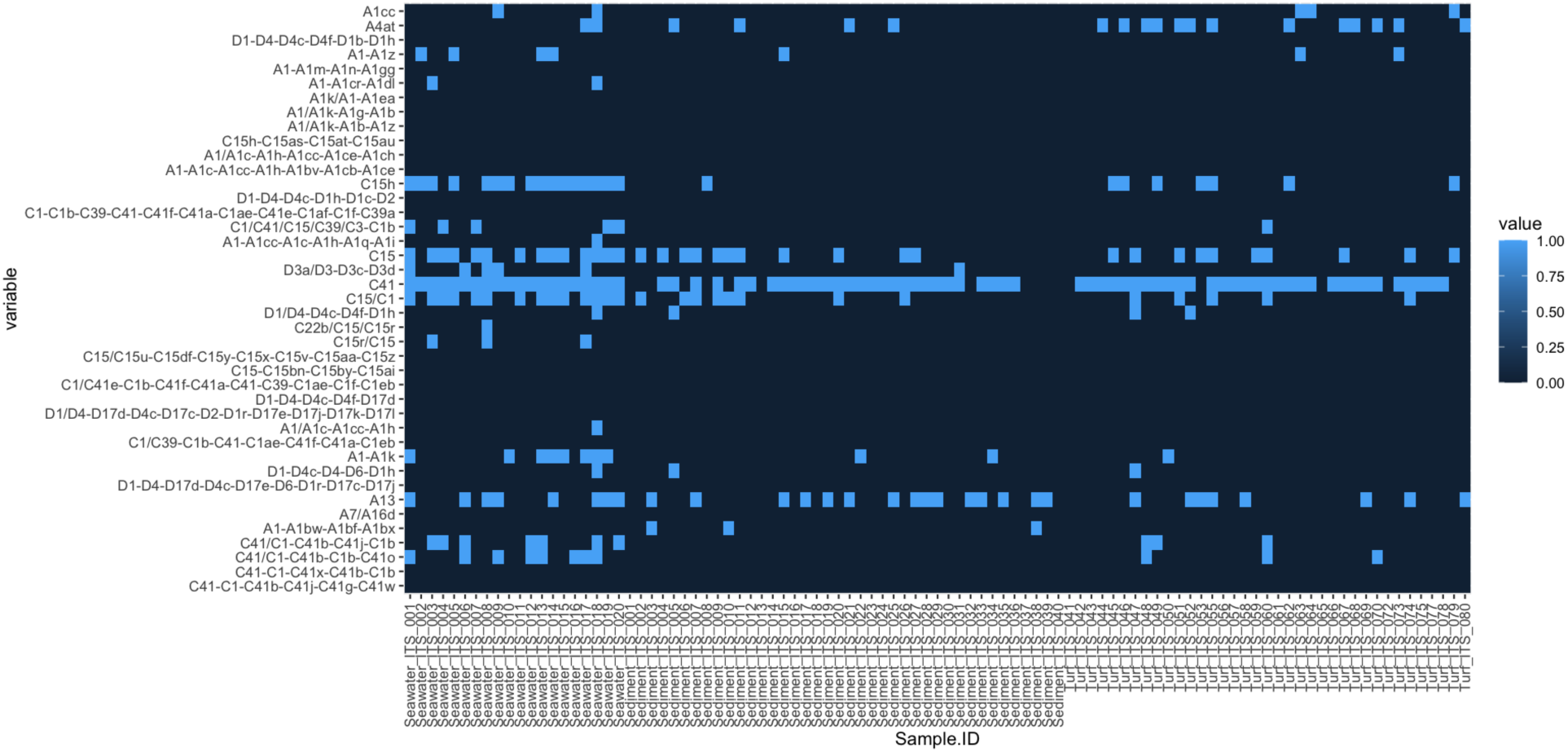
Presence/absence of complete DIV sets within environmental samples (columns) corresponding to the ITS2 type profiles (rows) found in coral samples. Light blue indicated all DIVs in the given ITS2 type profile were reported in the given sample. Dark blue indicates that at least one DIV was not present in that sample.

**Figure S7:**
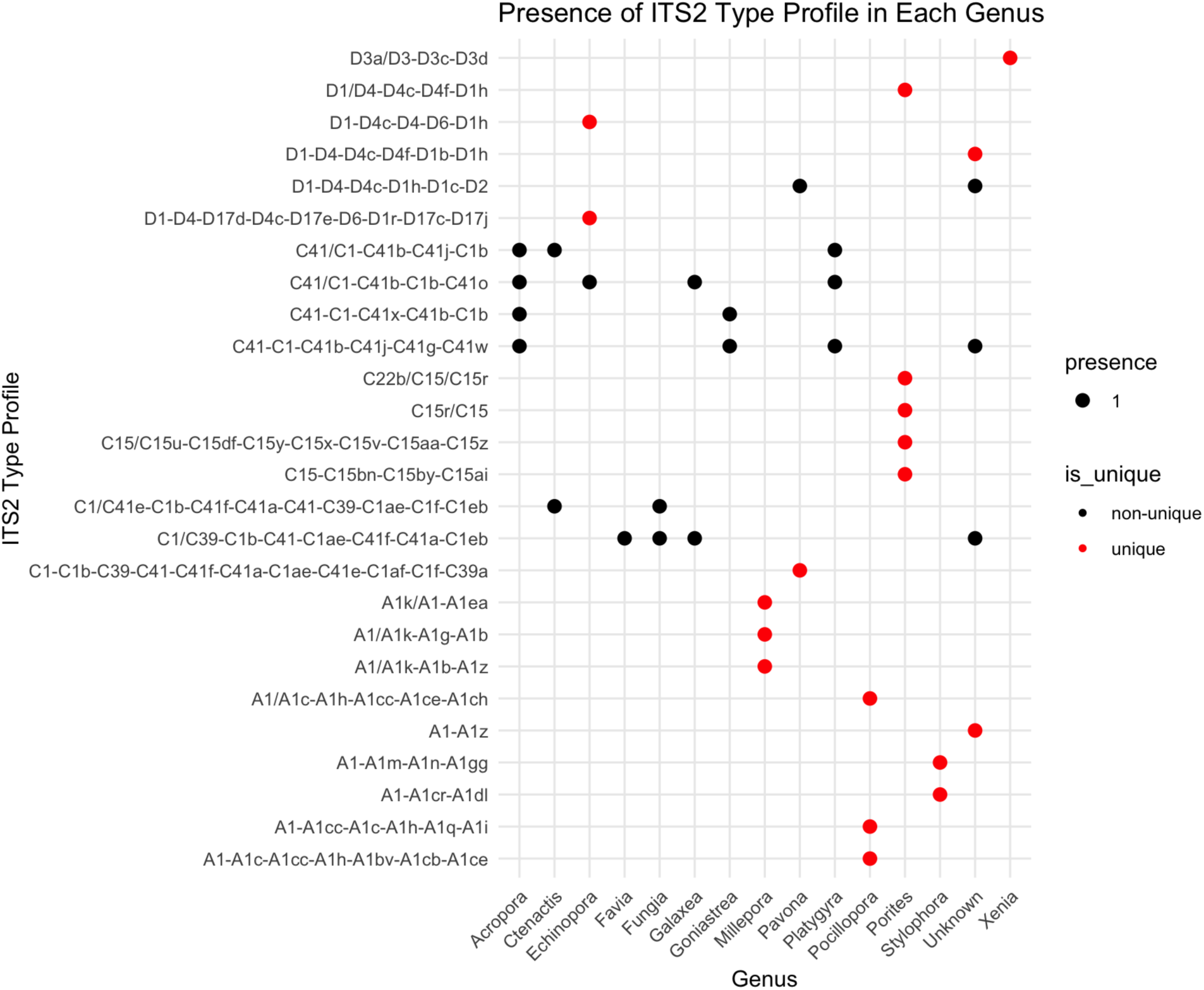
Presence of dominant ITS2 type profiles (>50% relative abundance within a sample, n=26) across coral genera assessed here. Both tissue and mucus samples are included for each genus - allowing genera with a single coral colony sampled (e.g. *Pavona*) to return two different dominant ITS2 type profiles. Red points show ITS2 type profiles that are unique to a single genus while black points show ITS2 type profiles present across genera.

**Figure S8:**
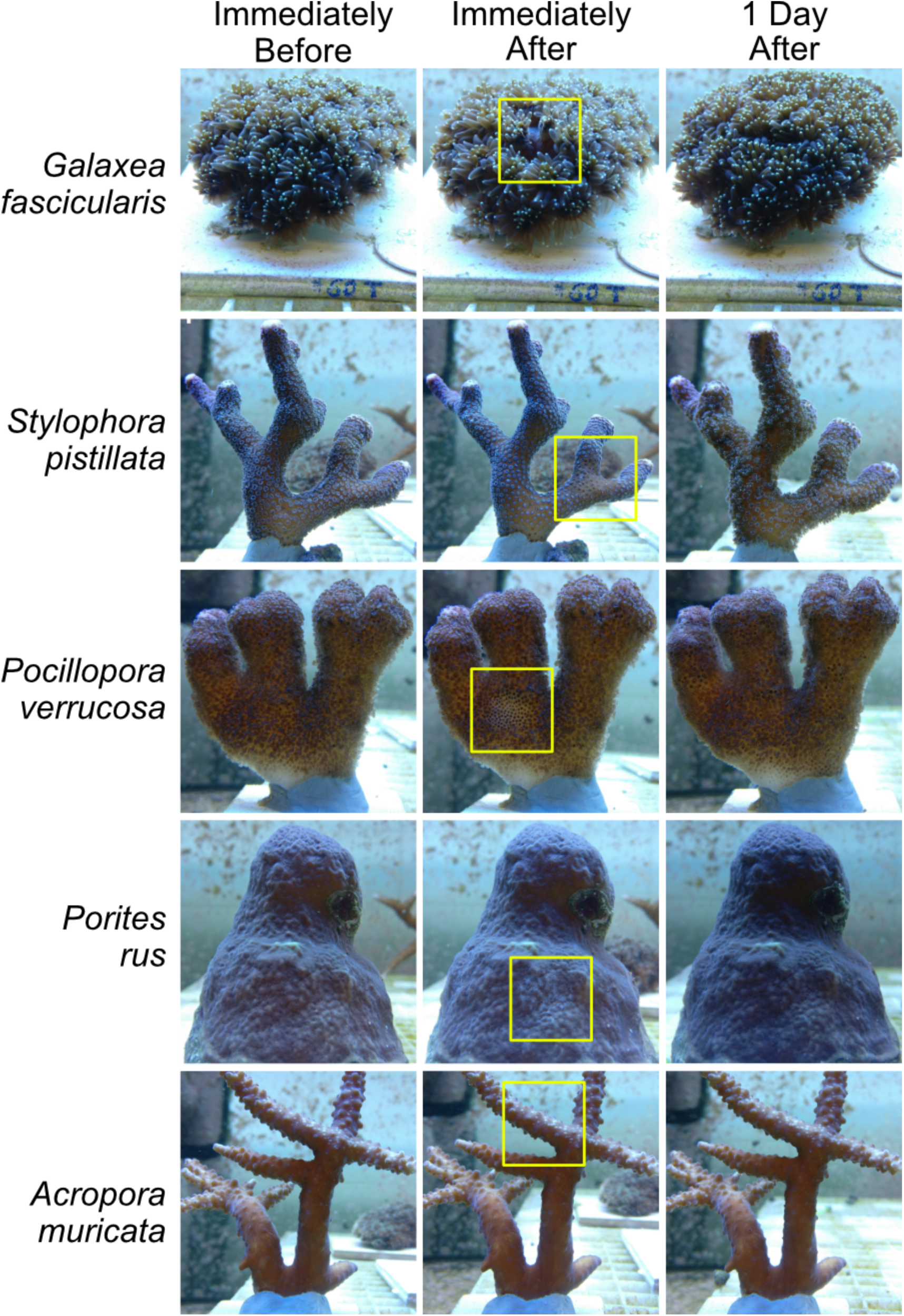
Resepentative colonies of each of 5 coral species subjected to simulated mucus sampling and their condition prior to, immediately after, and one day post sampling. The sampled area on each colony is demarcated with a yellow square.

